# Endocannabinoid and psychological responses to acute resistance exercise in trained and untrained adults

**DOI:** 10.1101/2023.09.08.556874

**Authors:** Zoe Sirotiak, Brandon T. Gallagher, Courtney A. Smith-Hernandez, Lucas J. Showman, Cecilia J. Hillard, Angelique G. Brellenthin

## Abstract

**Objective:** This study examined the effects of acute resistance exercise on circulating endocannabinoid (eCB) and mood responses in trained and untrained healthy adults. Methods: Thirty-two healthy adults (22.1 ± 2.9 years) were recruited from trained (reporting resistance exercise at least twice per week for ≥ previous three months) and untrained (performing no resistance exercise for ≥ previous three months) groups. Participants completed three sets of resistance exercise (16 repetitions at 50% 1-repetition max, 12 repetitions at 70% 1-repetition max, 8 repetitions at 80% 1-repetition max). Mood states, affect, and circulating eCB concentrations were evaluated before and after resistance exercise.

**Results:** There were significant decreases in AEA, PEA, and OEA levels following acute resistance exercise (p <0.05), with no significant group differences or group by time interactions. 2-AG did not change significantly. Positive affect increased significantly following resistance exercise (p =0.009), while negative affect decreased (p <0.001). Depression, anger, confusion, and total mood disturbance decreased significantly (p <0.05), while vigor increased significantly following resistance exercise (p =0.005). There were no significant group differences or group by time interactions for any psychological outcomes.

**Conclusion:** These results indicate that acute resistance exercise may reduce eCB and related lipid concentrations, which is opposite to the increase in lipids typically observed with acute aerobic exercise. Furthermore, psychological improvements occur after resistance exercise regardless of decreases in eCBs, supporting the notion that psychological changes with exercise likely occur through a wide variety of biological and environmental mechanisms.

## Introduction

The endocannabinoid (eCB) system is a widespread neuromodulatory network that regulates the function of numerous physiological processes (1). Critical components of this system include circulating eCBs, *N*-arachidonoylethanolamine (anandamide or AEA) and 2-arachidonoylglycerol (2-AG), which are endogenous ligands that bind to cannabinoid receptors and regulate neurotransmitter release at the local synapse (2,3). The eCB system is heavily represented in areas of the brain responsible for reward, cognition, and memory, and increased eCB activity has been associated with various psychological outcomes such as anxiety and depression (4,5). The eCB system is also activated by acute aerobic exercise, and numerous studies have indicated that eCBs may partially underlie the acute psychological benefits of aerobic exercise on mood and affect (6–8).

There have been multiple studies investigating the relationship between circulating eCBs and aerobic exercise. In general, an acute bout of moderate-to-vigorous intensity aerobic exercise reliably increases AEA in healthy adults, while very low or very high intensities do not (9–11).

Evidence is currently equivocal as to whether 2-AG increases in response to acute aerobic exercise (8–10). Furthermore, habitual levels of aerobic physical activity may also affect circulating concentrations of eCBs (7,8,12,13). Some studies have found that more active individuals have lower eCB activity at rest, whereas other studies have found higher eCB activity in active individuals (7,8,12,13). Differences in these findings may be the result of the sample characteristics (e.g., “exercise dependent” individuals, women with obesity). Nonetheless, growing evidence clearly suggests that the homeostatic functioning of the eCB system may be regulated by acute and chronic aerobic exercise, which could have therapeutic potential for conditions that are both improved by exercise and wherein the eCB system is implicated in the pathology (e.g., major depressive disorder, posttraumatic stress disorder, diabetes, obesity) (2,14,15). Despite the strong evidence associating aerobic exercise and the eCB system, there have been few investigations on the effects of resistance exercise, such as weightlifting, on eCBs. A single session of acute resistance exercise has been associated with activation of the eCB system in rats (16). In humans, acute resistance exercise has also been associated with significant increases in AEA and OEA concentrations, though PEA and 2-AG levels did not change significantly (17). Furthermore, a 15-week resistance exercise program was associated with significantly increased AEA levels in individuals with fibromyalgia but not healthy controls, suggesting that chronic resistance exercise training may be associated with changes in the eCB system in some conditions (18), although more research is needed.

Like aerobic exercise, acute resistance exercise has been shown to improve mood in both healthy and clinical groups (19,20). Chronic resistance exercise training has also been shown to yield similar improvements in mental health as aerobic exercise in patients with clinical anxiety or depression (19,20). However, there are currently no studies examining the involvement of the eCB system in these beneficial psychological effects. Since moderate-to-vigorous intensity aerobic exercise increases circulating eCBs, but low and high intensities do not, it is unclear whether resistance exercise, which traditionally involves short periods of high-intensity activity interspersed with periods of rest, would increase eCBs. However, skeletal muscle tissue, which is the primary tissue involved in resistance exercise, is thought to be a major site of eCB production in the periphery. Once they enter peripheral circulation, eCBs can travel across the blood-brain barrier to exert central effects (21,22). Furthermore, the eCB system is a highly plastic, stress-responsive, and adaptive system, and resistance exercise may be a reliable intervention for producing robust and rapid beneficial changes in muscle tissue and energy metabolism (2,14,15).

Thus, the eCB system is a plausible candidate mechanism underlying the effects of acute and chronic resistance exercise. Therefore, the purpose of this study was to determine the effects of acute resistance exercise on circulating concentrations of AEA and 2-AG and psychological outcomes in resistance-trained and untrained (i.e., novice) healthy young adults. We hypothesized that resistance exercise would increase eCBs and improve mood and affect and that these effects would be greater in the resistance trained rather than untrained individuals.

## Materials and Methods

### Study Participants

A total of 32 young adults aged 18-34 were recruited from the local community between October 2021 and June 2023. Prior to beginning the study, potential participants completed a screening process to assess eligibility status. Inclusion criteria included having performed resistance exercise at least twice per week for at least the past three months or performing no resistance exercise during this period. Participants who reported performing resistance exercise at least twice per week for at least the past three months were considered “trained” while those not performing any resistance exercise in the past three months were considered “untrained.” Exclusion criteria included having musculoskeletal problems that may be worsened by resistance exercise, previous fainting with blood draws, being pregnant or planning to become pregnant, being diagnosed with or taking prescription medication for chronic conditions including depression or anxiety, as well as using cannabis, tobacco, or other nicotine-based products. All research procedures were approved by the Institutional Review Board and participants provided written informed consent.

### Questionnaires

Physical Activity. Self-reported physical activity was measured through a 4-item version of the International Physical Activity Questionnaire – Short Form (IPAQ-SF) (23). Participants responded to questions about the frequency and session duration of moderate- and vigorous-intensity aerobic physical activities, resistance physical activities, as well as sitting over the past three months.

Affect. Positive and negative affect were measured using the Positive and Negative Affect Schedule (PANAS) (24). The 20-item measure asks about feelings and emotions over the past week, utilizing a Likert scale from 1 (very slightly or not at all) to 5 (extremely). The positive and negative affect subscales range from 10 to 50, with higher scores indicating higher levels of positive or negative affect. Reported internal consistency of the scale is adequate (positive affect subscale = α= 0.89, negative affect subscale α = 0.85) (24).

Anxiety. Trait anxiety was measured using the State-Trait Anxiety Inventory for Adults Form (STAI-T) (25). The 20-item measure asks about feelings in current moment, using a Likert scale from 1 (almost never) to 4 (almost always). Potential scores on the trait anxiety subscale of the measure ranges from 20 to 80, with norms and severity categories established. Higher scores indicate higher levels of trait anxiety. Reported internal consistency is adequate (α > 0.85) (25).

Mood states. The Profile of Mood States (POMS) measures six distinct mood states, resulting in the subscales tension, depression, anger, fatigue, confusion, and vigor (26). The 65-item measure asks about feelings during the past week and is rated on a 5-point Likert scale ranging from 0 (not at all) to 4 (extremely). A total mood disturbance measure can be calculated by adding the tension, depression, anger, fatigue, and confusion subscores and subtracting the vigor subscale, for a total possible score range of -24 to 177 with higher scores indicating higher levels of mood disturbance. Reported internal consistencies have been adequate (α > 0.70) (27).

### Laboratory Procedures

Each participant completed two laboratory visits at approximately the same time of day, separated by at least 7 days. Participants were asked to avoid vigorous-intensity exercise or resistance exercise during the 48 hours prior to each session to avoid influencing exercise performance and eCB levels. In addition, the participants were asked to avoid eating within 2 hours of each session.

### Familiarization Session

During the first session, participants provided written informed consent and answered questions to assess readiness to complete the session including general state of health, exercise in the previous 48 hours, substance use in the previous 24 hours, current illness, sleep, and use of over-the-counter medications. Participants then completed the IPAQ-SF, STAI, and PANAS measures. Peripheral blood pressure was measured automatically (Omron HEM-907xl) and was taken three times separated by 2 minutes. Height and weight were recorded, and body mass index (BMI) was calculated. Next, estimated 1-repetition maximum (1RM) testing on six weight machines (leg press, leg curl, leg extension, chest press, lat pulldown, and abdominal curl) was performed to set appropriate loads for the following session. A load was selected that the participant could lift anywhere between 1-10 repetitions, but no more, with proper form. A percentage of 1RM was calculated for each repetition from 1-10 based on the Landers equation, and an estimated 1RM was extrapolated for each machine based on the number of repetitions completed at the given load (28).

### Testing Session

The testing session began with the same questions to assess readiness to complete the session. Three peripheral blood pressures were taken, followed by a weight measurement.

Participants completed the PANAS and POMS, and then provided a blood sample via standard venipuncture to assess eCB concentrations prior to exercise. Resistance exercise was then performed on each of the six machines, with weights determined by 1RM results from the previous session. The first set consisted of 16 repetitions at 50% 1RM; the second set was 12 repetitions at 70% 1 RM; and the third set was 8 repetitions at 80% 1RM with two minutes of recovery between each set and machine. Immediately following exercise, another blood sample was collected. The PANAS and POMS questionnaires were completed again, followed by three peripheral blood pressure measurements.

### Blood Draw

Circulating concentrations of AEA and 2-AG were assessed using electrospray ionization, liquid chromatography/mass spectrometry (LC-MS/MS; Agilent Technologies, Santa Clara, CA), as detailed in previous literature (29,30). In addition, eCB-related ligands, palmitoylethanolamide (PEA) and N-oleoylethanolamine (OEA) were also assessed. These lipids are synthesized and released alongside AEA although they are not considered eCBs since they do not bind to CB1 or CB2 receptors. However, they are involved in appetitive, nociceptive, and inflammatory pathways that may also be stimulated by resistance exercise (31,32).

### Statistical methods

Lipid and psychological outcomes were analyzed using 2 (training group) x 2 (pre-, post-exercise) repeated measures analysis of variance (ANOVA). Endocannabinoid levels were adjusted through a natural logarithm transformation to better fit assumptions of equal variance and normality of the data. In an exploratory analysis, associations between changes in each lipid and changes in psychological outcomes, specifically total mood disturbance, positive affect, and negative affect, were analyzed using Spearman’s rank correlation with a Bonferonni adjustment for multiple comparisons. A p-value <0.05 was considered to be significant, and analyses were performed using SPSS Version 27.0 (33).

## Results

### Participant characteristics

Thirty-two participants completed the study, including 16 in the “trained” group and 16 in the “untrained” group. Participants were 59.4% women, with a mean age of 22.1 ± 2.9 years. There were no baseline group differences (p >0.05) in age, sex, blood pressure, or BMI. The trained group reported more moderate-to-vigorous aerobic physical activity (MVPA) and had higher 1RMs for all six machines (p <0.05; table 1).

**Table 1:** Baseline characteristics of untrained and trained groups.

|  | Untrained (n = 16) | Trained (n = 16) | Overall (n = 32) |
| --- | --- | --- | --- |
| Age (yr) | 22.56 (3.29) | 21.69 (2.52) | 22.13 (2.92) |
| Sex (male, female) | 5m, 11f | 8m, 8f | 13m, 19f |
| BMI (kg*m<sup>2</sup>) | 24.06 (3.21) | 24.83 (3.10) | 24.46 (3.12) |
| MVPA (min-wk<sup>-1</sup>; median [IQR]) | 300.00 (150.00-460.00)* | 715.00 (495.00-892.00) | 480.00 (260.00-810.00) |
| Trait Anxiety | 44.07 (7.60) | 40.63 (7.09) | 42.29 (7.43) |
| Positive Affect | 32.07 (5.46) | 35.44 (5.33) | 33.81 (5.57) |
| Negative Affect | 18.20 (7.21) | 18.94 (6.02) | 18.58 (6.52) |
| 1RM Leg Press (kg) | 287.25 (95.65)* | 508.75 (208.95) | 398.00 (195.48) |
| 1RM Leg Extension (kg) | 112.75 (22.13)* | 171.19 (63.73) | 141.97 (55.53) |
| 1RM Leg Curl (kg) | 105.94 (34.33)* | 145.69 (50.07) | 125.81 (46.81) |
| 1RM Chest Press (kg) | 87.25 (29.46)* | 155.19 (74.63) | 121.22 (65.62) |
| 1RM Lat Pulldown (kg) | 115.13 (33.33)* | 176.69 (74.11) | 145.91 (64.60) |
| 1RM Abdominal Curl (kg) | 57.94 (23.12)* | 83.31 (31.80) | 70.63 (30.23) |

Data presented are mean (SD) unless otherwise noted; BMI = body mass index; MVPA = moderate-vigorous physical activity; IQR=interquartile range; \*indicates significant difference from trained group ( $p < 0.05$ )

### Endocannabinoid concentrations

As shown in Figure 1, there were no differences in baseline concentrations of AEA, PEA, and OEA (p > 0.05), but trained participants had higher 2-AG than untrained participants (p = 0.044). AEA (F_1,30_ = 4.74, p = 0.038, Cohen’s *d =* -0.39), OEA (F_1,30_ = 13.13, p = 0.001, *d =* -0.48), and PEA (F_1,30_ = 7.15, p = 0.012, *d =* -0.65) decreased after resistance exercise in both groups. There were no changes in 2-AG (F_1,30_ = 0.25, p = 0.622, *d =* 0.09). There were no significant group differences or group by time interactions for changes in lipids.

**Fig 1:**
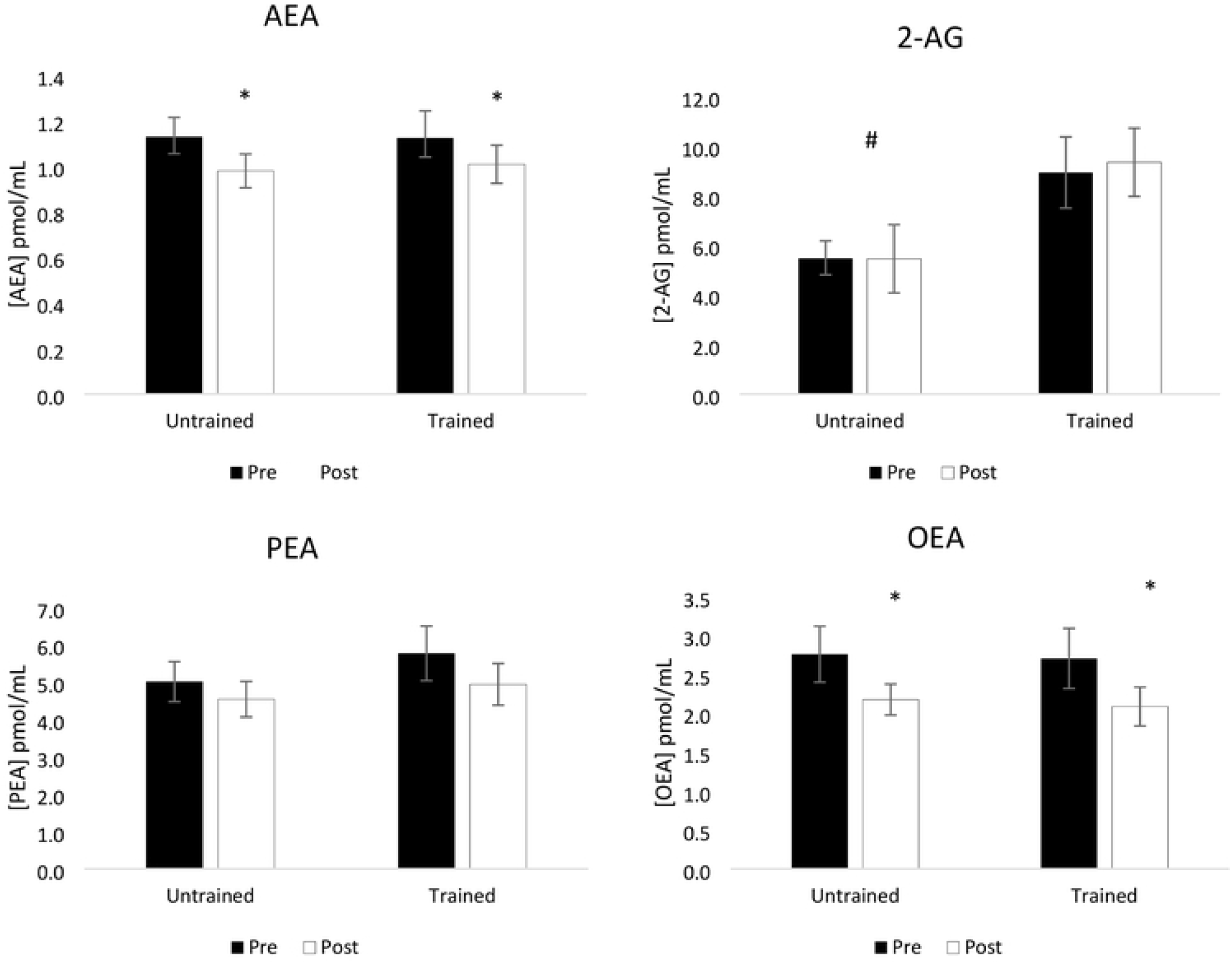
Lipid concentrations before and after acute resistance exercise.

Means and standard errors of AEA, 2-AG, PEA, and OEA before and after resistance exercise in the untrained and trained groups.

*Indicates significant (p <0.05) time effect for AEA, PEA, and OEA. # There was also a significant baseline group difference in 2-AG.

### Psychological variables

As shown in Table 2, negative affect decreased following resistance exercise (F_1,30_ = 14.99, p < 0.001), while positive affect increased (F_1,30_ = 7.80, p = 0.009). Total mood disturbance decreased following resistance exercise (F_1,30_ = 7.91, p = 0.009). Of the POMS subscales, confusion (F_1,30_ = 7.96, p = 0.008), anger (F_1,30_ = 8.67, p = 0.006), and depression (F_1,30_ = 4.80, p = 0.036) decreased significantly, while vigor increased significantly (F_1,30_ = 9.36, p = 0.005). Tension (F_1,30_ = 0.05, p = 0.828) and fatigue (F_1,30_ = 1.39, p = 0.248) did not change with resistance exercise. There were no significant group differences or group by time interactions for changes in psychological variables. Furthermore, there were no statistically significant correlations between changes in any lipid and changes in total mood disturbance, positive affect, or negative affect (all ps >0.14).

**Table 2:** Mood states and affect from before and after acute resistance exercise.

| | Mean change (SD) | p-value for time effect | $\eta^2$ for time effect |
| --- | --- | --- | --- |
| **Tension** |  | 0.828 | 0.00 |
| Untrained | 0.00 (3.60) |  |  |
| Trained | -0.31 (4.41) |  |  |
| **Depression** |  | 0.036 | 0.14 |
| Untrained | -0.25 (2.29) |  |  |
| Trained | -2.94 (5.35) |  |  |
| **Anger** |  | 0.006 | 0.22 |
| Untrained | -0.75 (1.39) |  |  |
| Trained | -2.13 (3.65) |  |  |
| **Vigor** |  | 0.005 | 0.24 |
| Untrained | 2.75 (5.04) |  |  |
| Trained | 2.19 (4.04) |  |  |
| **Fatigue** |  | 0.248 | 0.04 |
| Untrained | 1.81 (2.26) |  |  |
| Trained | -0.56 (3.60) |  |  |
| **Confusion** |  | 0.008 | 0.21 |
| Untrained | -1.00 (2.53) |  |  |
| Trained | -1.56 (2.61) |  |  |
| **Total Mood Disturbance** |  | 0.009 | 0.21 |
| Untrained | -2.94 (7.46) |  |  |
| Trained | -9.69 (16.34) |  |  |
| **Positive Affect** |  | 0.009 | 0.21 |
| Untrained | 3.81 (5.56) |  |  |
| Trained | 1.63 (5.45) |  |  |
| **Negative Affect** |  | < 0.001 | 0.33 |
| Untrained | -1.19 (2.34) |  |  |
| Trained | -1.88 (2.13) |  |  |

There were no significant group differences or group by time interactions for any of the outcomes reported.

## Discussion

The results of this study indicate that resistance exercise reduced circulating concentrations of AEA, PEA, and OEA. This result is contrary to existing acute aerobic exercise-related literature showing increases in circulating eCBs, specifically AEA, with moderate-to-vigorous intensity aerobic exercise (11,34). This finding is also contrary to the limited resistance exercise literature, which shows that acute resistance exercise is associated with increased AEA concentrations (17), although there were substantial differences in the acute resistance exercise protocols used in that study and the present study (e.g., fasted vs. not fasted; acute resistance exercise protocol of 1RM testing on 2 machines vs. 3 sets on 6 machines, respectively). There was no significant change in 2-AG, which is consistent with some aerobic and resistance exercise studies (9,17,34). Decreased eCBs were not expected based on aerobic exercise literature to date, and the finding indicates that aerobic and resistance training may have different acute effects on the eCB system. One possible explanation for this difference may be the intensity of the exercise performed affecting the energy systems involved. Most aerobic exercise studies investigating eCB responses to date consist of continuous, moderate-to-vigorous intensity aerobic challenges (≥30 minutes) primarily stressing the aerobic energy system (8,10,11,35,36). While the average rating of perceived exertion across all sets of the resistance exercise in this study was also of a moderate-to-vigorous intensity (untrained RPE=15.0[1.1]; trained RPE=14.8 [1.2]; p=0.61), the intermittent design of the resistance exercise protocol likely challenged the phosphocreatine and anaerobic energy systems (36), which may influence the response of the eCB and other modulatory systems. The eCB system may be activated to a greater extent by aerobic exercise due to the unique effects of aerobic metabolism in the periphery. Specifically, OEA acts on the peroxisome proliferator-activated receptor-alpha (PPAR-alpha), which is also involved in aerobic metabolism (34,37). However, acute isometric exercise usually in one limb, which is primarily anaerobic, has also been found to increase circulating AEA (17,38). Thus, whole body resistance exercise, despite involving anaerobic pathways, may have additional unique effects on the eCB system.

This is the first study to show decreases in eCBs in response to acute resistance exercise, so follow-up studies to confirm these results are clearly warranted. Given the medium to large effect sizes for decreases in PEA and OEA, we achieved 75% and 95% power, respectively, to detect a true underlying effect. Although our achieved power for decreases in AEA was smaller at 57%, AEA, PEA, and OEA are synthesized along the same pathway (39), providing physiological support for our similar findings among these three lipids. While this study is unable to explain the reduction in lipids, there may be some possible reasons why decreases in these select lipids may be beneficial. For example, AEA is involved in analgesia and 2-AG is involved in psychological effects (18,40). OEA is involved in suppressing appetite, and finally, PEA is associated with anti-inflammatory effects (31,32). Given the adaptations that must occur after resistance exercise, reduced lipid signaling in light of these pathways may relay important information regarding muscle pain, appetite stimulation, and increased inflammation that often occur as a result of resistance exercise. Somewhat surprisingly, though, resistance exercise training status did not influence acute changes in eCBs. Some aerobic exercise literature suggests that habitual levels of aerobic exercise or levels of cardiorespiratory fitness (often used as a proxy for recent engagement in aerobic exercise), reduce basal eCB activity yet also promote more robust changes in response to acute exercise (8,13). Since the eCB system is a homeostatic system, its ability to respond quickly to stressors and return to resting levels thereafter may represent better functioning of the system at large. Similar decreases in eCBs after resistance exercise in the trained and untrained groups in this study may suggest that training status does not affect eCB response to acute resistance exercise.

Despite reductions or no changes in eCBs after resistance exercise, there were significant improvements in affect and mood in both groups. There were also no correlations between changes in eCBs and changes in psychological outcomes. These findings indicate that circulating concentrations of eCBs may not be a major component of psychological responses to resistance exercise, although more research is needed to support this finding. Regulation of other systems, such as endogenous opioids, brain-derived neurotrophic factor, hypothalamic-pituitary-adrenal axis, or inflammatory pathways could be investigated as alternative explanations (41,42).

Considering non-physiological causes such as the training environment or staff-participant interactions are important as well. With increased awareness of placebo effects in psychological outcomes of exercise, the placebo effect must be considered as a possible contributor to our findings, too (43).

This study has limitations. This study recruited healthy young adults, and the results of this study may not apply to other populations. Similarly, participants with diagnosed psychiatric conditions such as anxiety and depression disorders or those taking medications for these conditions were not eligible for this study, and therefore effects on mood or affect cannot be generalized to clinical psychiatric populations or those taking psychiatric medications. Inclusion criteria for the trained group required at least three months of resistance training, but it is possible that a longer period of training may be needed to see group-based effects. Relatedly, this study was also cross-sectional, and both aerobic and resistance exercise training trials are critically needed to explore the long-term effects of exercise training on the eCB system and downstream physical and mental health outcomes. Finally, effects were only measured once immediately before and once immediately after resistance exercise performance, and it is unclear how eCB and psychological outcomes may change beyond the time points measured or during exercise itself.

## Conclusions

This study showed decreases in circulating AEA, PEA and OEA concentrations following acute resistance exercise. There were no changes in 2-AG. There were significant increases in positive affect and decreases in negative affect and several negative mood states after exercise. This finding suggests that unlike with aerobic exercise, acute mood and affect improvements following resistance exercise may occur via alternative psychobiological or environmental (i.e., context and setting) mechanisms. Future studies are warranted to confirm and expand upon these initial findings in larger and more diverse healthy and clinical samples.

## Acknowledgements

We acknowledge the W.M. Keck Metabolomics Research Laboratory (Office of Biotechnology, Iowa State University, Ames, IA) for providing analytical instrumentation.

